# Analytical Performance of a Standardized Kit for Mass Spectrometry-based Measurements of Human Glycosaminoglycans

**DOI:** 10.1101/2021.02.04.429694

**Authors:** Davide Tamburro, Sinisa Bratulic, Souad Abou Shameh, Nikul K Soni, Andrea Bacconi, Francesca Maccari, Fabio Galeotti, Karin Mattsson, Nicola Volpi, Jens Nielsen, Francesco Gatto

## Abstract

Glycosaminoglycans (GAGs) are long linear sulfated polysaccharides implicated in processes linked to disease development such as mucopolysaccharidosis, respiratory failure, cancer, and viral infections, thereby serving as potential biomarkers. A successful clinical translation of GAGs as biomarkers depends on the availability of standardized GAG measurements. However, owing to the analytical complexity associated with the quantification of GAG concentration and structural composition, a standardized method to simultaneously measure multiple GAGs is missing. In this study, we sought to characterize the analytical performance of a ultra-high-performance liquid chromatography coupled with triple-quadrupole tandem mass spectrometry (UHPLC-MS/MS)-based kit for the quantification of 17 GAG disaccharides. The kit showed acceptable linearity, selectivity and specificity, accuracy and precision, and analyte stability in the absolute quantification of 15 GAG disaccharides. In native human samples, here using urine as a reference matrix, the analytical performance of the kit was acceptable for the quantification of CS disaccharides. Intra- and inter-laboratory tests performed in an external laboratory demonstrated robust reproducibility of GAG measurements showing that the kit was acceptably standardized. In conclusion, these results indicated that the UHPLC-MS/MS kit was standardized for the simultaneous measurement of GAG disaccharides allowing for comparability of measurements and enabling translational research.

**Summary:** Analytical performance of a kit for standardized GAG measurements, based on an established UHPLC-MS/MS method

## 1 Introduction

Glycosaminoglycans (GAGs) are a family of long linear polysaccharides consisting of repeating disaccharide units (*1*). Different structural disaccharides of GAGs have been characterized. In humans, the most prevalent classes are chondroitine sulfate (CS) [(→ 3)-β-D- GalNAc(1→ 4)-β-D-GlcA or α-L- IdoA(1→], heparan sulfate (HS) [(→ 4)-α-D-GlcNAc or α-D- GlcNS(1→ 4)-β-D-GlcA or α-L-IdoA (1→], and hyaluronic acid (HA) [(→ 3)-β-D-GlcNAc(1 → 4)-β-D-GlcA(1→] where GalNAc is N-acetylgalactosamine, GlcA is glucuronic acid, IdoA is iduronic acid, GlcNAc is N-acetylglucosamine, and GlcNS is N-sulfoglucosamine. CS and HS disaccharides can each be further modified with O-sulfo groups in up to three positions. The modifications afford eight different sulfation patterns that modulate the biophysical properties of CS and HS in the extracellular matrix.

This physio-chemical diversity of GAG disaccharides enables highly diverse biological functions and implicates GAGs in health- and disease-relevant processes such as cell proliferation and wound healing (*2, 3*). For this reason, GAGs showed promise as biomarkers in several diseases, like mucopolysaccharidosis (*4*), respiratory failure (*5, 6*), cancer (*7–9*), and viral infections (*10*).

Precise measurements of GAG concentration and structural composition at the level of individual GAG disaccharides are crucial for structure-function studies on the importance of GAGs in human health and disease. Moreover, standardization of GAG measurements is imperative for future clinical translation in biomarker discovery. In recent years, ultra-high-performance liquid chromatography coupled with triple-quadrupole tandem mass spectrometry (UHPLC-MS/MS) emerged as the reference platform for robust and reliable GAG quantification (*11–23*). However, no standardized kit is currently available hampering the comparability of findings and translational research.

Here, we describe a kit based on a previously established method for GAG extraction and detection by Volpi et al. (*12*). In short, this method relies on the enzymatic depolymerization of GAGs into disaccharides and their subsequent derivatization using 2-aminoacridone (AMAC). Separation is achieved using ultra-high-performance liquid chromatography (UHPLC) and detection using electrospray ionization triple-quadrupole mass spectrometry (ESI-MS/MS) through multiple reaction monitoring (MRM) analysis. Even though this method has been described extensively in the literature in different versions (*12, 18*), a standardized kit based on it is lacking and its analytical performance characteristics are unknown. Thus, we sought to perform a systematic evaluation of the kit’s analytical performance characteristics.

## 2 Material and Methods

### 2.1 Glycosaminoglycan quantification method

We independently quantified concentrations (in µg mL^-1^) of 8 CS disaccharides (0s CS, 2s CS, 6s CS, 4s CS, 2s6s CS, 2s4s CS, 4s6s CS, Tris CS), 8 HS disaccharides (0s HS, 2s HS, 6s HS, Ns HS, Ns6s HS, Ns2s HS, 2s6s HS, Tris HS) and HA. Also, we calculated the total concentration of CS and HS as the sum of the corresponding disaccharide concentrations. Unless otherwise specified, we performed GAG extraction, detection, and quantification in a single commercial laboratory (reference laboratory, Lablytica Life Science AB, Uppsala, Sweden) using a single lot of Elypta MIRAM™ Glycosaminoglycan Kit.

The method was performed following Elypta MIRAM™ Glycosaminoglycan Kit instructions for use. All reagents and consumables used were contained in the kit. This method was based on a previously established protocol for glycosaminoglycan (GAG) extraction and detection by Volpi et al. (*12*) (Figure 1A). Briefly, the method consisted of an enzymatic digestion assay using *Chondroitinase ABC* and *Heparinase I-II-III* to depolymerize GAGs in the sample into disaccharides. Disaccharides were subsequently labeled using 2-aminoacridone. The samples were then injected into an ultra-high-performance liquid chromatography (UHPLC) coupled with electrospray ionization triple-quadrupole mass spectrometry system (ESI-MS/MS, Waters® Acquity I-class Plus Xevo TQ-S micro) for disaccharide separation and detection. The peaks of the 17 disaccharides were acquired at pre-specified retention times across six transitions using multiple reaction monitoring (MRM) analysis implemented in the mass spectrometry software (Waters® TargetLynx). We used the mass spectrometry software (Waters® TargetLynx) for peak integration, construction of calibration curves, and quantification. We exported the results processed data in Excel format and imported it into *R* (4.0.2) for secondary analysis.

**Figure 1.**
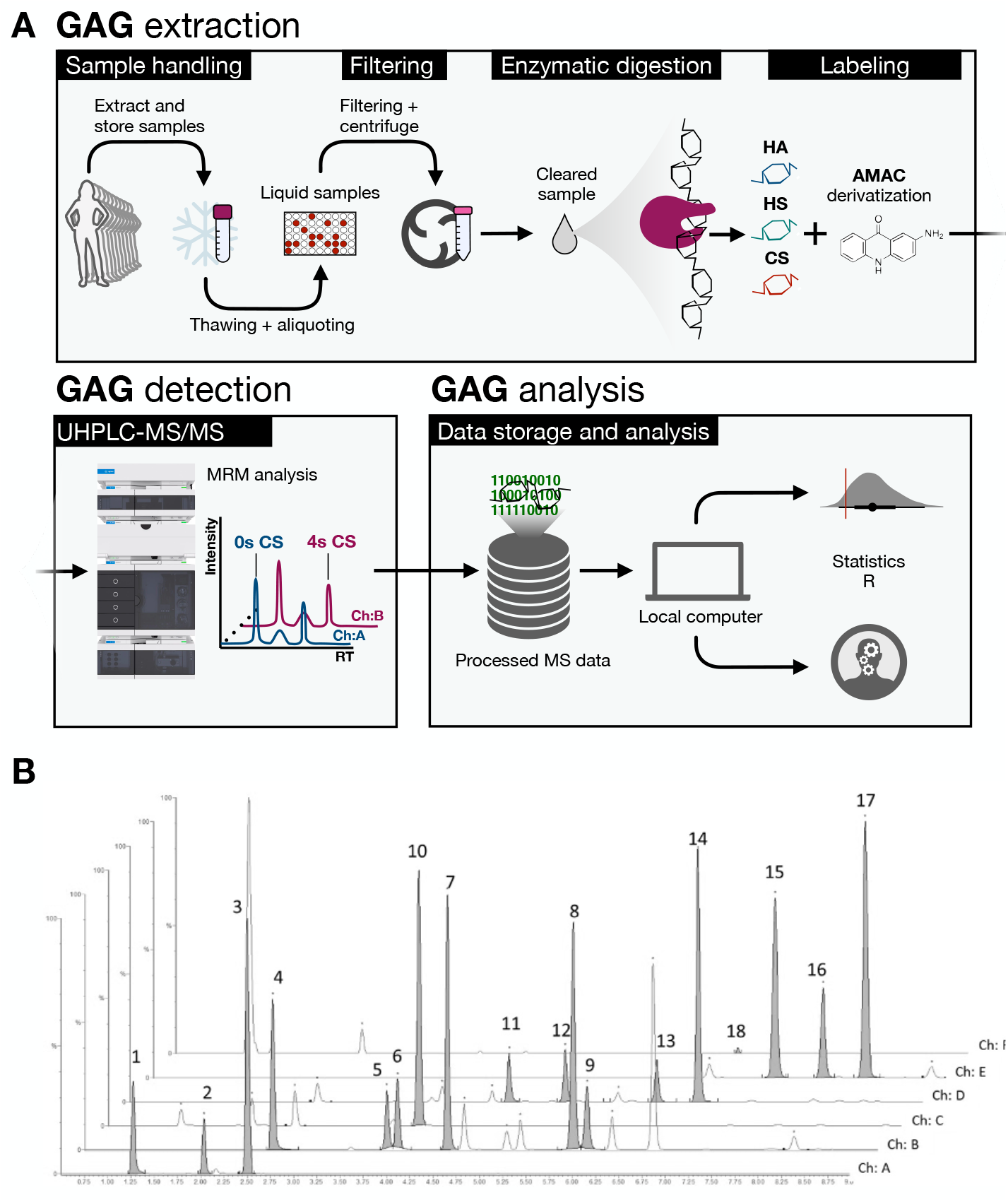
A) Overview of the method described in Volpi et al. (*12*) on which the UHPLC-MS/MS kit is based on. B) A representative chromatogram for a mixture of 17 purified disaccharides. The 17 disaccharide peaks are acquired in one multiple reaction monitoring (MRM) run across six different channels (Ch: A to F) based on mass transitions associated with analytes. Key - 1: Tris H; 2: Ns6s HS, 3: Ns2S HS, 4: Tris CS; 5: 2s4S CS; 6: 2s6s HS, 7: 2s6s CS, 8: 2s HS, 9: 2s CS, 10: Ns HS, 11: 4s6s CS, 12: 6s HS, 13: 4s CS, 14: 6s CS, 15: 0s HS, 16: HA, 17: 0s CS.

### 2.2 Analytical performance tests

We carried out a battery of tests to characterize the linearity, selectivity, specificity, accuracy, precision of the calibrators, carryover, stability of the calibrators as well as linearity, selectivity, specificity, accuracy, precision of the calibrators, stability, recovery, and matrix effects intra- and inter-laboratory precision in native samples (see SI Methods for detailed experimental descriptions). The tests were designed using recommendations from the following CLSI Approved Guidelines: C62-A (LC-MS Methods), EP05-A3 (Precision), EP06-A (Linearity), EP17-A2 (Detection Capability), EP25-A (Stability), C50-A (MS General Principles) C24 (Statistical Quality Control). We prespecified acceptance criteria for all tests except for matrix effects (see SI Methods for pre-specified acceptance criteria.

The tests were conducted either on blank samples, standard samples, or “proxy urine” and native urine samples, as appropriate. For standard samples, we created a set of three standard GAG solutions at three different concentrations (low, medium, and high). We prepared the standard GAG solution at the “high” level by mixing the highest level of the calibrator sample for all disaccharides in milli-Q water. Next, we serially diluted the sample at the “high” level to “medium” and “low” levels (1: 0.50: 0.25 v/v) using Milli-Q water. For proxy urine samples, we prepared a “proxy urine” pool by mixing urine collected from healthy donors. Next, we depleted GAGs from the proxy urine pool by recovering the filtrate resulting from ultracentrifugation (14000 *g* at 9 °C for 60 minutes) in the filter provided in the kit. We spiked the standard GAG solution into the proxy urine at the beginning of sample preparation (before filtration) or immediately after filtration during sample preparation. For native urine samples, we sourced native urine from self-rated healthy adult donors and collected them in a polypropylene jar at room temperature. We kept samples frozen (- 80 °C) until the analysis. The study was approved by the Ethical Committee (Etikprövningsmyndigheten) in Gothenburg, Sweden on February 8, 2018 (#737-17). Further details on samples are provided in the SI Methods.

## 3 Results

### 3.1 Overview of the standardized kit method

The kit – based on a previously described method illustrated in Figure 1A (*12*) - consisted of a instructions-for-use protocol, reagents, and consumable for sample preparation to extract GAGs from frozen samples and subsequent MRM analysis for GAG detection and quantification using a UHPLC-MS/MS method. Seventeen disaccharides (8 for CS, 8 for HS, and HA) – here on referred to as analytes – could be independently and simultaneously measured with the kit. The kit included the standard solution and instructions to prepare a calibration curve for each analyte starting from the provided highest calibration level (the nominal concentration of the disaccharide at each calibration level was provided in Table S1); a mixture of 17 purified disaccharides (one per analyte) to aid the operator in the adjustment of eventual drifts in retention times, and four quality control (QC) samples to be used in every MS run to monitor inter-sequence variability. In Figure 1B, representative chromatograms are illustrated for the afore-mentioned mixture of 17 purified disaccharides quantified using the kit (see Figure S1 for chromatograms derived from the standard GAG solution).

We characterized the analytical performance of the kit in terms of calibration capability and performance in native human samples.

### 3.2 Characterization of the kit analytical performance: calibration

First, we characterized the calibration curve parameters: linearity, detection capability, selectivity and specificity, accuracy and precision, carry-over, and disaccharide stability in the auto-sampler. This process established the performance of the calibration curve over a specific range for each disaccharide.

In the second part, we performed the characterization of the kit in terms of GAGs extraction, detection, and quantification in native (human) samples by measuring the following parameters: selectivity and specificity, recovery, matrix effect, linearity response, accuracy and precision, and disaccharide stability.

#### 3.2.1 Linearity and detection capability of the calibrators

We tested the linearity of the calibration curve for each disaccharide. We prepared nine levels of calibration of each disaccharide in triplicates, injecting each replicate two times. We pre-specified acceptance criteria for a level to be included in the final calibration curve in terms of acceptable coefficient of variation (CV), which was required to be lower than 25% (30% for the lowest level), and deviation with the respect to the nominal concentration after back-calculation, which was required to be lower than 25%. We defined the upper limit of quantification (ULoQ) and low limit of quantification (LLoQ) for each disaccharide as the highest and lowest calibration levels meeting the acceptance criteria, respectively. We defined the range the linearity as the concentration between LLoQ and ULoQ.

In Table 1, we reported for each disaccharide the number of calibration levels within the range of linearity between LLoQ and ULoQ, including the coefficient of determination (R^2^) of the calibration curve and the coefficient of variation (CV) and deviation of back-calculated concentration with respect to the nominal concentration at the LLoQ and ULoQ.

**Table 1.** Linearity of the calibration curve for each disaccharide including the number of calibrator levels, coefficient of determination (R^2^), and nominal concentration, coefficient of variation (CV), and deviation from back-calculated concentration for the lower limit of quantification (LLoQ) and upper limit of quantification (ULoQ). Acceptable values are marked in bold. Key: N.C. – not calibrated.

| Disaccharide | N levels | $R^2$ | LLoQ<br>[$\mu\text{g mL}^{-1}$] | LLoQ<br>CV | LLoQ<br>deviation | ULoQ<br>[$\mu\text{g mL}^{-1}$] | ULo<br>CV | ULoQ<br>deviation |
| --- | --- | --- | --- | --- | --- | --- | --- | --- |
| **0s CS** | 8 | 1.000 | 0.10 | 13% | -14% | 5.33 | 2% | -1% |
| **4s CS** | 7 | 0.996 | 1.42 | 7 | -19 | 42.87 | 2 | 2 |
| **6s CS** | 5 | 0.980 | 1.15 | 4 | 1 | 11.15 | 2 | -10 |
| **4s6s CS** | 4 | 0.999 | 0.06 | 12 | -6 | 0.31 | 8 | -1 |
| **2s4s CS** | 3 | 0.999 | 0.04 | 15 | 4 | 0.11 | 11 | 1 |
| **2s6s CS** | 6 | 0.999 | 0.04 | 31 | 6 | 0.60 | 5 | 1 |
| **Tris CS** | 0 | N.C. | N.C. | N.C. | N.C. | N.C. | N.C. | N.C. |
| **2s CS** | 2 | 1.000 | 0.02 | 22 | 0 | 0.04 | 14 | 0 |
| **HA** | 6 | 0.997 | 3.51 | 5 | 23 | 60.00 | 2 | 3 |
| **Tris HS** | 2 | 1.000 | 0.24 | 21 | 0 | 0.43 | 39 | 0 |
| **Ns2s HS** | 3 | 0.999 | 0.46 | 26 | 2 | 1.44 | 20 | 0 |
| **2s6s HS** | 0 | N.C. | N.C. | N.C. | N.C. | N.C. | N.C. | N.C. |
| **Ns6s HS** | 4 | 0.998 | 0.37 | 22 | 9 | 2.03 | 6 | 1 |
| **Ns HS** | 6 | 0.996 | 0.48 | 10 | -16 | 8.20 | 6 | -4 |
| **0s HS** | 6 | 0.998 | 2.70 | 6 | -14 | 46.27 | 1 | -3 |
| **2s HS** | 3 | 0.995 | 0.02 | 21 | -7 | 0.06 | 14 | -1 |
| **6s HS** | 7 | 0.999 | 0.09 | 14 | -29 | 2.66 | 7 | -2 |

For 15 of 17 disaccharides (all except Tris CS and 2s6s HS), we obtained a calibration curve with acceptable linearity and detection capability. In cases where the calibration curve could not be constructed (for Tris CS and 2s6s HS), GAG quantification relied on the ratio between the observed peak area and the corresponding peak area at the highest level of the provided calibrator.

#### 3.2.2 Selectivity and specificity of the calibrators

We tested the selectivity and specificity to each disaccharide by inspecting the presence of peaks of an area greater than 20 % LLoQ for that disaccharide in blank samples. For each disaccharide, no peak could be detected at the expected retention time for that disaccharide in any of the blank samples. Selectivity and specificity for the kit were therefore deemed acceptable for all disaccharides.

#### 3.2.3 Accuracy and precision of the calibrators

We tested the accuracy and precision of the calibration curves by creating a set of three standard GAG solutions at three different concentrations (low, medium, and high). In Table 2 and 3, we reported the accuracy (percentage difference between nominal concentration for a disaccharide in the standard GAG solution and measured concentration) over a 2-day experiment and the precision (as CV) for each disaccharide at each standard GAG solution concentration (low, medium, and high).

**Table 2.** Accuracy (in terms of % deviation from nominal concentration in the standard GAG solution) at three concentration levels on two separate days. Acceptable values are marked in bold. Key: N.D. – not detected.

| Disaccharide | Low |  | Medium |  | High |  |
| --- | --- | --- | --- | --- | --- | --- |
|  | Day 1 | Day 2 | Day 1 | Day 2 | Day 1 | Day 2 |
| **0s CS** | -61% | -59% | -20% | -17% | -9% | -5% |
| **4s CS** | 14 | 16 | -7 | -5 | -4 | -2 |
| **6s CS** | -23 | -20 | 0 | 3 | -1 | 4 |
| **4s6s CS** | 1 | -3 | 3 | 0 | 2 | 3 |
| **2s4s CS** | N.D. | N.D. | -4 | 1 | -6 | 5 |
| **2s6s CS** | 1 | -5 | 1 | 1 | 2 | 4 |
| **Tris CS** | N.D. | N.D. | N.D. | N.D. | N.D. | N.D. |
| **2s CS** | N.D. | N.D. | N.D. | N.D. | 100 | 79 |
| **HA** | -27 | -22 | -12 | -9 | -3 | 0 |
| **Tris HS** | N.D. | N.D. | N.D. | N.D. | 7 | 6 |
| **Ns2s HS** | N.D. | N.D. | 37 | -25 | 10 | -41 |
| **2s6s HS** | N.D. | N.D. | N.D. | N.D. | N.D. | N.D. |
| **Ns6s HS** | 26 | 8 | 19 | 4 | 12 | -2 |
| **Ns HS** | -18 | -31 | -1 | -14 | 0 | -12 |
| **0s HS** | 5 | 1 | 6 | 0 | -3 | -7 |
| **2s HS** | N.D. | N.D. | 28 | N.D. | 8 | -49 |
| **6s HS** | 4 | -29 | 13 | -21 | 3 | -30 |

**Table 3.** Precision (in terms of CV% in the estimated concentration for each disaccharide in the standard GAG solution) at three concentration levels over two days. Acceptable values are marked in bold. Key: N.D. – not detected.

| **Disaccharide** | **Low** |  | **Medium** |  | **High** |  |
| --- | --- | --- | --- | --- | --- | --- |
|  | Day 1 | Day 2 | Day 1 | Day 2 | Day 1 | Day 2 |
| **0s CS** | 2% | 3% | 4% | 4% | 3% | 5% |
| **4s CS** | 2 | 3 | 1 | 2 | 3 | 3 |
| **6s CS** | 1 | 3 | 1 | 3 | 3 | 5 |
| **4s6s CS** | 13 | 12 | 8 | 10 | 7 | 7 |
| **2s4s CS** | N.D. | N.D. | 15 | 17 | 8 | 14 |
| **2s6s CS** | 12 | 13 | 6 | 6 | 8 | 8 |
| **Tris CS** | N.D. | N.D. | N.D. | N.D. | N.D. | N.D. |
| **2s CS** | N.D. | N.D. | N.D. | N.D. | 224 | 416 |
| **HA** | 1 | 4 | 2 | 3 | 3 | 4 |
| **Tris HS** | N.D. | N.D. | N.D. | N.D. | 20 | 22 |
| **Ns2s HS** | N.D. | N.D. | 36 | 58 | 22 | 40 |
| **2s6s HS** | N.D. | N.D. | N.D. | N.D. | N.D. | N.D. |
| **Ns6s HS** | 16 | 24 | 16 | 20 | 14 | 18 |
| **Ns HS** | 5 | 11 | 5 | 12 | 3 | 11 |
| **0s HS** | 2 | 5 | 3 | 6 | 3 | 5 |
| **2s HS** | N.D. | N.D. | 20 | N.D. | 18 | 41 |
| **6s HS** | 8 | 26 | 9 | 29 | 7 | 26 |

The accuracy was acceptable for virtually all detected CS and HA disaccharides at all 3 concentrations on both Day 1 and Day 2, as well as for most detected HS disaccharides at medium and high concentrations on Day 1 (but not on Day 2). Similarly, both intraday and inter-day precision were acceptable for all detected CS and HA disaccharides at all 3 concentrations as well as for intraday precision for most detected HS disaccharides at all 3 concentrations. We attributed the results reported in Tables 2 and 3 that did not meet the acceptance criteria to poor signal acquisition.

#### 3.2.4 Carry-over in the calibrators

We tested the impact of the carry-over by inspecting the presence of peaks of an area greater than 20 % LLoQ for that disaccharide in blank samples immediately after the acquisition of the highest calibration curve level. For each disaccharide, no peak could be detected at the expected retention time for that disaccharide in any of the blank samples. Carry-over was therefore considered negligible for all disaccharides.

#### 3.2.5 Disaccharide stability in the autosampler

We tested the stability of disaccharides stored in the autosampler at 10 °C by monitoring the peak area of each analyte over 14 days at the third-highest calibration curve level for that disaccharide. In Table S2, we reported the change in peak areas for a given disaccharide at a given time point relative to the corresponding peak area at the initial time point.

The stability in the auto-sampler of all disaccharides except Ns2s HS was acceptable over 6 days (at least 5 of 6-time points with <30% deviation from Day 0). We found that all disaccharides except 4s CS had poor stability after 14 days in the autosampler.

### 3.3 Characterization of the kit analytical performance: concentration estimation

Having established the performance of the calibration curve for each disaccharide, we could estimate disaccharide concentrations using the so-characterized calibration curves. In the second part, we, therefore, performed the characterization of the kit in terms of estimation of the disaccharide concentration in native samples by measuring the following performance parameters: selectivity and specificity, recovery, matrix effect, linearity response, accuracy, and precision and disaccharide stability. For the sake of consistency, we chose human urine as the reference matrix for the native samples.

#### 3.3.1 Recovery

We tested recovery by first generating a GAG-depleted sample (referred to as proxy urine, SI Methods); and next by spiking the proxy urine with a set of three standard GAG solutions at three different concentrations (low, medium, and high) either at the beginning of the sample preparation or immediately after filtration during sample preparation. In Table S3, we reported the recovery of each disaccharide after filtration across five replicates.

The results indicated acceptable recovery (<25% deviation) for all detectable CS disaccharides at all three concentration levels (except 2s CS, acceptable only at the highest level); for all detectable HS disaccharides at the “low” and “medium” concentration levels (except for 2s HS); and for HA at the “medium” concentration level.

#### 3.3.2 Matrix effect

We tested matrix effects to evaluate how endogenous compounds in native urine interfered with the measurement of the analytes. We spiked proxy urine from 6 healthy donors and milli-Q water samples in triplicates with the above-described set of standard GAG solutions at two concentration levels, “high” and low”. In Table S4, we reported the matrix effect as the ratio (in %) between the disaccharide concentration in 6 proxy urine samples versus the milli-Q water sample at the two concentration levels, as well as the average matrix effect in proxy urine.

The matrix effect was moderate (between 42% and 89%) in CS disaccharides at the “high” level and low (78% to 103%) at the “low” level, indicative of signal suppression due to matrix effects. We found moderate-to-high matrix effects in HA and HS disaccharides at the “high” level and moderate-to-low at the “low” level.

#### 3.3.3 Accuracy and precision in native samples

We tested the accuracy of disaccharide concentrations in native samples by spiking the proxy with the same set of standard GAG solutions as described above at three different concentration levels (low, medium, and high). In Table 4, we reported the accuracy in the concentration of each CS disaccharide in a three-day experiment by two operators on a given analysis day. We computed the intraday precision in terms of coefficient of variation from the same data and reported in Table 5. We then computed the intraday precision as the coefficient of variation from the same data and reported in Table 5.

**Table 4.** Accuracy of disaccharide concentration in proxy urine sample spiked at three concentration levels of a standard GAG solution over a three-day experiment by two operators. Note that accuracy for HA and HS disaccharides could not be reliably estimated in this experiment and it was omitted. Acceptable values marked in bold. Key: N.D. – not detected.

| Disaccharide | Low |  |  | Medium |  |  | High |  |  |
| --- | --- | --- | --- | --- | --- | --- | --- | --- | --- |
|  | Day 1 | Day 2 | Day 3 | Day 1 | Day 2 | Day 3 | Day 1 | Day 2 | Day 3 |
| **0s CS** | 21% | 20% | 21% | 31% | 34% | 31% | 32% | 34% | 39% |
| **4s CS** | 10 | 0 | 6 | 19 | 15 | 15 | 5 | 5 | 4 |
| **6s CS** | 5 | -12 | -9 | 21 | 9 | 11 | 15 | 13 | 20 |
| **4s6s CS** | 22 | 19 | 19 | 31 | 30 | 27 | 26 | 29 | 30 |
| **2s4s CS** | N.D. | N.D. | N.D. | N.D. | N.D. | N.D. | 28 | 30 | 31 |
| **2s6s CS** | 13 | 8 | 11 | 26 | 27 | 26 | 20 | 22 | 27 |
| **Tris CS** | N.D. | N.D. | N.D. | N.D. | N.D. | N.D. | N.D. | N.D. | N.D. |
| **2s CS** | N.D. | N.D. | N.D. | N.D. | N.D. | N.D. | N.D. | N.D. | N.D. |

**Table 5.** Precision (in terms of CV%) of disaccharide concentration in proxy urine sample spiked at three concentration levels of a standard GAG solution over a three-day experiment by two operators. Note that precision for HA and HS disaccharides could not be reliably estimated in this experiment and it was omitted. Acceptable values marked in bold. Key: N.D. – not detected.

| **Disaccharide** | **Low** |  |  | **Medium** |  |  | **High** |  |  |
| --- | --- | --- | --- | --- | --- | --- | --- | --- | --- |
|  | Day 1 | Day 2 | Day 3 | Day 1 | Day 2 | Day 3 | Day 1 | Day 2 | Day 3 |
| **0s CS** | 5% | 7% | 6% | 3% | 13% | 12% | 4% | 6% | 17% |
| **4s CS** | 5 | 10 | 12 | 1 | 5 | 5 | 2 | 2 | 3 |
| **6s CS** | 6 | 16 | 15 | 2 | 14 | 12 | 4 | 4 | 14 |
| **4s6s CS** | 13 | 13 | 14 | 8 | 9 | 10 | 7 | 7 | 8 |
| **2s4s CS** | N.D. | N.D. | N.D. | N.D. | N.D. | N.D. | 13 | 12 | 14 |
| **2s6s CS** | 11 | 13 | 12 | 8 | 11 | 10 | 7 | 7 | 14 |
| **Tris CS** | N.D. | N.D. | N.D. | N.D. | N.D. | N.D. | N.D. | N.D. | N.D. |
| **2s CS** | N.D. | N.D. | N.D. | N.D. | N.D. | N.D. | N.D. | N.D. | N.D. |

We were unable to reliably determine the accuracy and precision for HA and HS disaccharides because their estimated concentrations were close to the respective LLoQ even for the “high” level standard GAG solutions. We attributed this partly to matrix effects as later discussed.

The accuracy in native samples was acceptable (<30% deviation from the nominal concentration) for all detectable CS disaccharides except for 0s CS, where the accuracy was below 70% in “medium” and “high” level samples - albeit never below 60%. The intraday precision in native samples was acceptable (CV < 25%) for all detectable CS disaccharides.

#### 3.3.4 Disaccharide stability in native samples

We tested the stability of each disaccharide in the native matrix over 14 days. Specifically, we prepared two proxy urine samples and spiked them with the above-described set of standard GAG solutions at two concentration levels (“high” and low”). We stored the spiked samples at -20 °C and measured disaccharide concentrations on day 1 and day 14. In Table S5, we reported the percentage difference in the disaccharide concentration at a given level (“high” or “low” at Day 1 and Day 14 compared to the corresponding nominal concentration.

The stability in native samples was acceptable (<30% deviation from the nominal concentration) for all detectable CS disaccharides on Day 1 and for 4 of 6 detectable CS disaccharides on Day 14, where the remaining two CS disaccharides (0s CS and 4s6s CS) deviated from the nominal concentrations between 33% and 37%, respectively. Note that the stability for HA and HS disaccharides could not be reliably estimated in this experiment and it was omitted.

#### 3.3.5 Selectivity and specificity in native samples

We tested selectivity and specificity to each disaccharide by inspecting the presence of peaks of an area greater than 20 % LLoQ in proxy urine - without any spiked GAG solution. For each disaccharide, no peak could be detected in proxy urine at the expected retention time for that disaccharide. Selectivity and specificity in native samples were therefore acceptable for all disaccharides.

#### 3.3.6 Linearity in native samples

We tested the linearity of disaccharide concentrations in native samples by spiking proxy with the set of standard GAG solutions as described above at nine different concentration levels. In Table 6, we reported the linearity for each disaccharide in terms of coefficient of determination (R^2^) for the linear regression between peak areas and concentration across the nine levels.

**Table 6.**
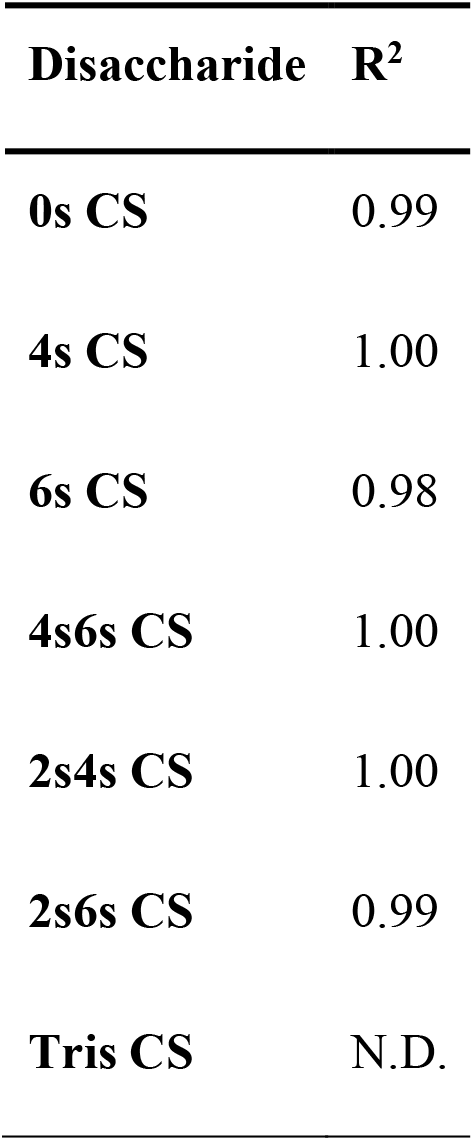
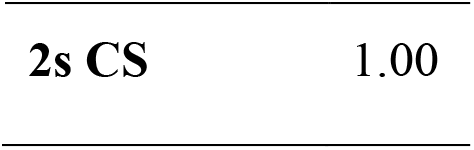
Linearity of disaccharides in proxy urine samples (in terms of coefficient of determination R^2^ between peak areas and concentration levels) of a standard GAG solution serially diluted to generate nine concentration levels. Note that stability for HA and HS disaccharides could not be reliably estimated in this experiment and it was omitted. Acceptable values marked in bold. Key: N.D. – not detected.

| Disaccharide | $R^2$ |
| --- | --- |
| **0s CS** | 0.99 |
| **4s CS** | 1.00 |
| **6s CS** | 0.98 |
| **4s6s CS** | 1.00 |
| **2s4s CS** | 1.00 |
| **2s6s CS** | 0.99 |
| **Tris CS** | N.D. |
| 2s CS | 1.00 |

The linearity was acceptable for all detectable CS disaccharides (R^2^ > 0.95). Note that we were unable to reliably estimate the linearity for HA and HS disaccharides.

### 3.4 External validation of kit analytical performance

We sought to validate the hereby presented analytical performance specifications by performing tests of intra-laboratory and interlaboratory precision in a GLP-compliant external laboratory (Lablytica Life Science AB, Uppsala, Sweden).

#### 3.4.1 Intra-laboratory precision

We tested intra-laboratory precision by monitoring the disaccharide concentration in four QC samples included in the kit throughout 14 independent experiments (runs). We focused on the two properties of the GAG profile, non-sulfated CS (0s CS) and total CS, namely the sum of all measured CS disaccharides in a sample. Two QC samples were synthetic samples spiked with standard GAG solutions at high or low concentration (∼14.5 µg mL^-1^ and ∼1.7 µg mL^-1^ for total CS, respectively). The other two QC samples were native samples with known high or low total GAG concentration (∼12.5 µg mL^-1^ and ∼4.2 µg mL^-1^ for total CS, respectively). In Figure 2, the estimated concentration of the two key GAG properties (total CS and 0s CS) were plotted across runs. The concentrations for all measured CS GAGs were shown in Figure S2. In Table S6, we reported the CV for each disaccharide in the 4 QC samples.

**Figure 2.**
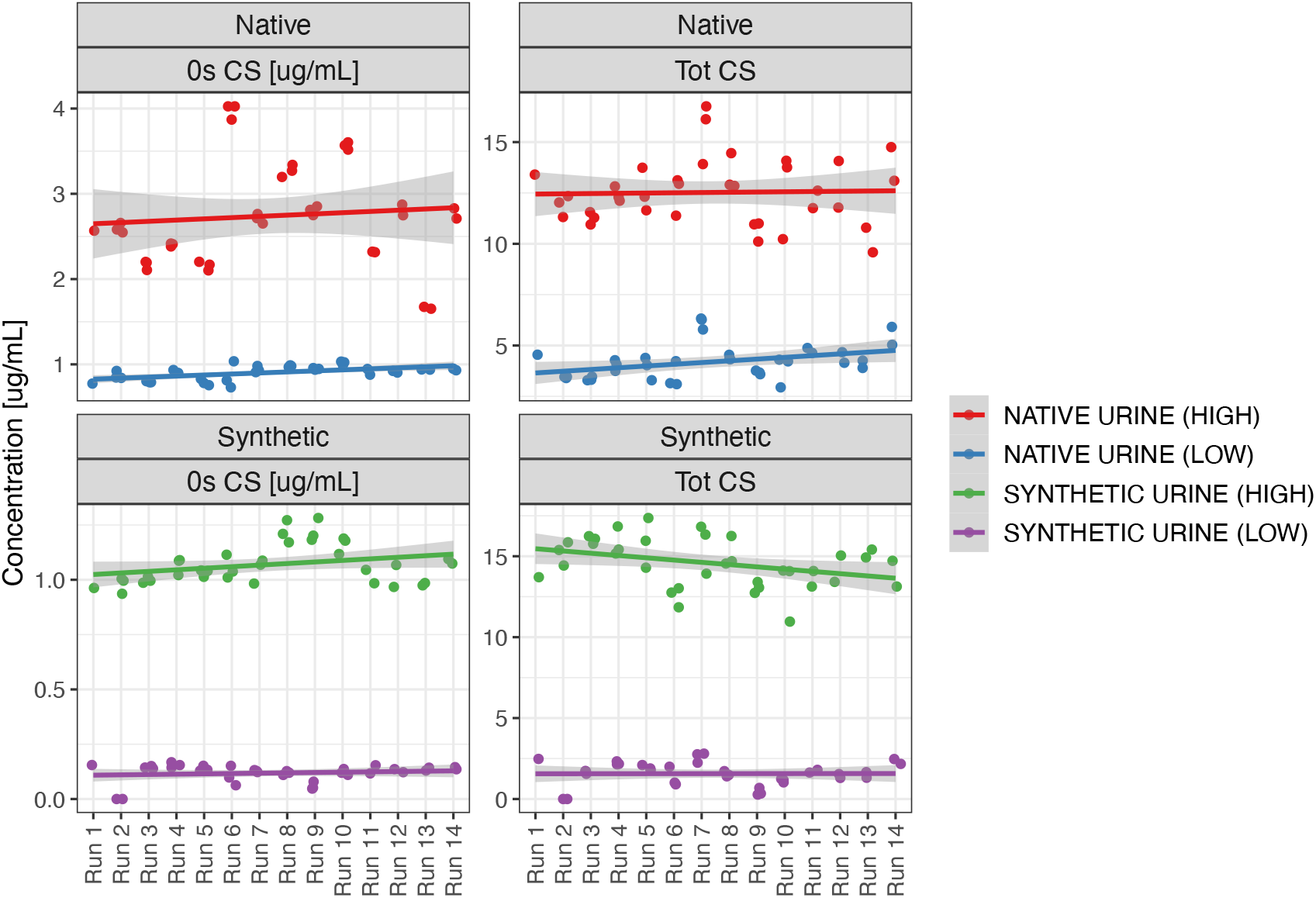
Total CS and 0s CS concentration (in µg mL^-1^) in 4 QC samples (in duplicates) across 14 runs in an external laboratory. The line represents the least square regression for disaccharide concentration across runs (shaded area represents 95% confidence interval on the regressed mean).

The intra-laboratory precision was acceptable for all major CS disaccharides. The precision for HS disaccharides was acceptable only in “low” concentration samples, while for HA it was acceptable only in synthetic QC samples. In general, the concentration for di-sulfated and tri-sulfated disaccharides as well as 2s disaccharides was below LLoQ in all QC samples and therefore precision estimates should be interpreted with caution.

#### 3.4.2 Inter-laboratory precision

We tested inter-laboratory precision by comparing the disaccharide concentration of two key GAG properties (total CS and 0s CS) in a panel of nine native urine samples from healthy donors independently analyzed in the reference laboratory versus the external laboratory. We found that the total CS and 0s CS concentration estimates from the reference laboratory versus the external laboratory, were strongly correlated (Pearson correlation coefficient *R* > 0.95) (Figure 3).

**Figure 3.**
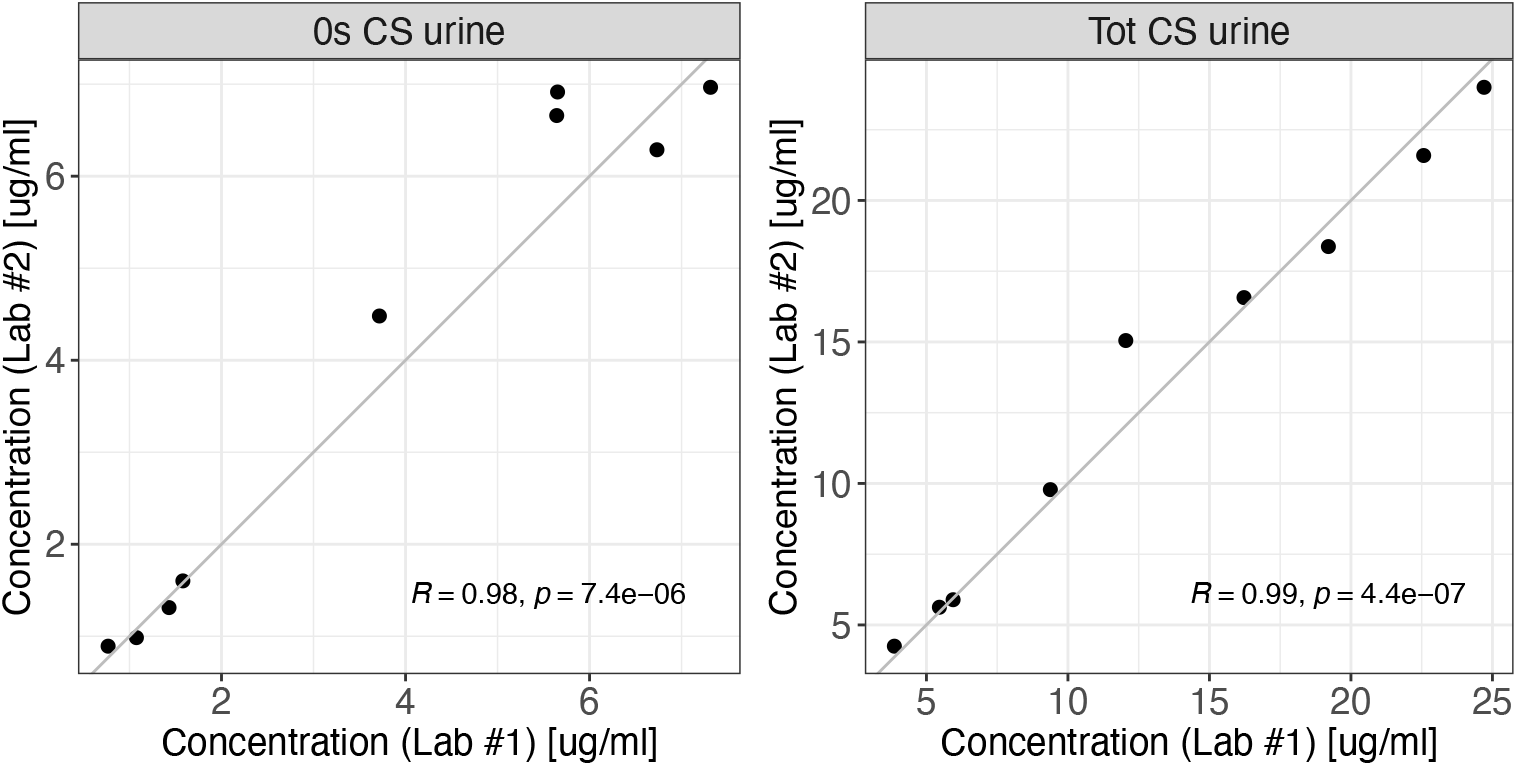
0s CS and total CS concentration in 9 native urine samples from healthy donors as estimated in the reference laboratory (Lab #1) versus the external laboratory (Lab #2). The diagonal line represents the identity line. The Pearson correlation coefficient *R* for the correlation is displayed with its *p*-value (permutation test).

## 4 Discussion

GAGs are increasingly recognized as key molecular actors in mechanisms central to human physiology and pathology such as structural support within the extracellular matrix and regulation of cell signaling (*2, 24*). These mechanisms ultimately control phenotypes that, in disease, can culminate in physical and mental disabilities in mucopolysaccharidosis (*4*), jeopardize wound healing (*25*), promote cancer (*26*), lead to respiratory failure (*3*), or enable binding of viruses such as SARS-CoV-2 (*10*). GAGs have therefore been proposed as potential biomarkers for these different diseases (*4, 27, 28*). We and others have demonstrated the added advantage of measuring GAGs non-invasively, for example in blood and urine (*5–8, 29*). However, moving beyond biomarker discovery to clinical translation necessitates the standardization of GAG measurements. UHPLC-MS/MS has emerged in the last decade as the gold standard for rapid quantification of GAG concentration and disaccharide composition. Many methods were published that illustrated efficient separation of up to 19 disaccharides in biological samples (*11–23*). However, none of these reported extensive analytical performance testing. Therefore, here we comprehensively characterized the analytical performance of a kit for GAG measurements using an UHPLC-MS/MS method which has been previously described (*12*).

Overall, the kit efficiently separated 17 disaccharides and exhibited excellent selectivity and specificity to all disaccharides with negligible carryover and sufficient stability for typical laboratory work shifts. The calibrators were accurate and precise for 15 of 17 disaccharides over a range of concentrations covering approximately one-order of magnitude for each disaccharide. Notably, most above-cited methods appeared to rely on a single point of calibration for all disaccharides interrogated, thereby returning a relative concentration of each disaccharide in the sample (i.e. a mass fraction composition). Exceptions to these are the methods presented by Tomatsu et al. (2014), which reported calibration curves for 7 disaccharides (16), and Yang et al. (2012), which calibration curves for the same 17 disaccharides as those detected here (23). Compared to published methods, the here-characterized kit for absolute quantification of GAGs adequately was found to be capable to calibrate as many as 15 disaccharides simultaneously.

In native samples, here using urine as the reference matrix, we demonstrated the robust and accurate analytical performance characteristics of the kit. Critically for analyses of biological and clinical samples, the kit enabled the quantification of CS disaccharides within a calibration range that captured physiological values spanning one order of magnitude. We were unable to validate the results on HA and HS disaccharides because the concentrations recovered in urine were below the LLoQ for virtually all of these disaccharides. This could simply reflect a low abundance of urinary HA and HS in physiological conditions compared to CS. The hypothesis is in line with previous reports in which the total HA and HS concentration in urine were measured as ∼20% of the total CS concentration, almost an order of magnitude less abundant (*8, 18*). Nevertheless, we could not rule out the hypothesis that the kit was underperforming in the quantification of HA and HS disaccharides in the urine. We attributed one possibility to the here-observed matrix effects, which showed moderate to strong peak area suppression in HA and HS in the urine. Another explanation could be a less efficient extraction yield during enzymatic digestion of HA and HS disaccharides as compared to CS disaccharides. Overall, we showed that the kit had an acceptable analytical performance for the quantification of CS disaccharides in native samples, while its performance in HA and HS warranted further investigation.

We deemed the kit standardized for CS measurements given that the intra-laboratory and inter- laboratory precision tests produced acceptable results across two independent laboratories. Specifically, the here-described kit proved capable to simultaneously calibrate 15 of 17 disaccharides with high linearity (R^2^ > 0.99 for all disaccharides except 6s CS wherein R^2^ = 0.98) and with high intra-laboratory precision using 14 replicates of four control urine samples (CV ranging 8 to 22% at “high” concentration and 8 to 42% at “low” concentration for the major CS disaccharides, here defined as >5% of total CS). In comparison, Tomatsu et al. (2014) reported a method that could quantify 7 disaccharides (2 keratan sulfate disaccharide, 3 HS, and 2 CS) with linearity R^2^ ranging 0.982 to 0.993 across two orders of magnitude and with an intra-assay precision CV ranging 2 to 15% using three control serum samples (16). Yang et al. (2012) proposed a method for the quantification of 17 disaccharides (same as those detected by the kit here described) with linearity R^2^ ranging 0.976 to 0.999 across one order of magnitude -but no estimates on assay precision (23). Wei et al. (2013) described a method that could quantify 12 HS disaccharides in relative concentrations (mass fraction %) with an intra-assay precision CV ranging 1 to 21% for the major HS disaccharides (>5% of total HS) using one control serum sample (17). Overall, the kit had performance characteristics comparable with previously developed methods while extending the breadth of GAG quantification to 15 disaccharides.

In conclusion, we verified the analytical performance of a kit for GAG disaccharide quantification using an LC/MS-MS method. The findings suggest that the kit had standardized characteristics for the absolute quantification of 15 disaccharides and precise and accurate quantification of CS disaccharides in native samples. Future work should be directed towards the inclusion of disaccharides here omitted, for example, keratan sulfate disaccharides, and to improve the analytical performance in the quantification of HA and HS disaccharides in the urine.

## 5 Acknowledgments

This work was supported by the Knut and Alice Wallenberg Foundation, Cancerfonden, and the IngaBritt och Arne Lundbergs Forskningsstiftelse (J.N.) and in part by the European Union’s Horizon 2020 research and innovation programme under grant agreement No 849251 (Fr.G.). The authors wish to thank Angelo Limeta for assistance with the figures.

## 6 Author contributions

D.T. designed the experimental plan. D.T., S.S., and N.K. performed the experiments. D.T., S.B., and Fr.G., analyzed and interpreted the data. F.M., Fa.G., A.B., K.M., N.V. provided critical feedback in the experimental design, experiment execution, and data interpretation. Fr.G. and J.N. conceived the study and provided funding. Fr.G. drafted the manuscript. All authors critically reviewed the manuscript in its final form.

## References

1. J. D. Esko, K. Kimata, U. Lindahl, in Essentials of Glycobiology, A. Varki, R. D. Cummings, J. D. Esko, H. H. Freeze, P. Stanley, C. R. Bertozzi, G. W. Hart, M. E. Etzler, Eds. (Cold Spring Harbor Laboratory Press, Cold Spring Harbor (NY), ed. 2nd, 2009; http://www.ncbi.nlm.nih.gov/books/NBK1900/).

2. N. Afratis, C. Gialeli, D. Nikitovic, T. Tsegenidis, E. Karousou, A. D. Theocharis, M. S. Pavão, G. N. Tzanakakis, N. K. Karamanos, Glycosaminoglycans: key players in cancer cell biology and treatment. The FEBS Journal. 279, 1177–1197 (2012).

3. A. B. Souza-Fernandes, P. Pelosi, P. R. Rocco, Bench-to-bedside review: The role of glycosaminoglycans in respiratory disease. Crit Care. 10, 237 (2006).

4. R. Lawrence, J. R. Brown, F. Lorey, P. I. Dickson, B. E. Crawford, J. D. Esko, Glycan-based biomarkers for mucopolysaccharidoses. Mol Genet Metab. 111, 73–83 (2014).

5. E. P. Schmidt, K. H. Overdier, X. Sun, L. Lin, X. Liu, Y. Yang, L. A. Ammons, T. D. Hiller, M. A. Suflita, Y. Yu, Y. Chen, F. Zhang, C. Cothren Burlew, C. L. Edelstein, I. S. Douglas, R. J. Linhardt, Urinary Glycosaminoglycans Predict Outcomes in Septic Shock and Acute Respiratory Distress Syndrome. Am J Respir Crit Care Med. 194, 439–449 (2016).

6. E. P. Schmidt, G. Li, L. Li, L. Fu, Y. Yang, K. H. Overdier, I. S. Douglas, R. J. Linhardt, The circulating glycosaminoglycan signature of respiratory failure in critically ill adults. J Biol Chem. 289, 8194–8202 (2014).

7. F. Gatto, K. A. Blum, S. S. Hosseini, M. Ghanaat, M. Kashan, F. Maccari, F. Galeotti, J. J. Hsieh, N. Volpi, A. A. Hakimi, J. Nielsen, Plasma Glycosaminoglycans as Diagnostic and Prognostic Biomarkers in Surgically Treated Renal Cell Carcinoma. European Urology Oncology. 1, 364–377 (2018).

8. F. Gatto, N. Volpi, H. Nilsson, I. Nookaew, M. Maruzzo, A. Roma, M. E. Johansson, U. Stierner, S. Lundstam, U. Basso, J. Nielsen, Glycosaminoglycan Profiling in Patients’ Plasma and Urine Predicts the Occurrence of Metastatic Clear Cell Renal Cell Carcinoma. Cell Rep. 15, 1822–1836 (2016).

9. T. M. Clausen, G. Kumar, E. K. Ibsen,M.S. Ørum-Madsen, A. Hurtado-Coll, T. Gustavsson,M.Ø. Agerbæk, F. Gatto, T. Todenhöfer, U. Basso, M. A. Knowles, M. Sanchez-Carbayo, A. Salanti, P. C. Black, M. Daugaard, A simple method for detecting oncofetal chondroitin sulfate glycosaminoglycans in bladder cancer urine. Cell Death Discovery. 6, 1–7 (2020).

10. T. M. Clausen, D. R. Sandoval, C. B. Spliid, J. Pihl, H. R. Perrett, C. D. Painter, A. Narayanan, S. A. Majowicz, E. M. Kwong, R. N. McVicar, B. E. Thacker, C. A. Glass, Z. Yang, J. L. Torres, G. J. Golden, P. L. Bartels, R. N. Porell, A. F. Garretson, L. Laubach, J. Feldman, X. Yin, Y. Pu, B. M. Hauser, T. M. Caradonna, B. P. Kellman, C. Martino, P. L. S. M. Gordts, S. K. Chanda, A. G. Schmidt, K. Godula, S. L. Leibel, J. Jose, K. D. Corbett, A. B. Ward, A. F. Carlin, J. D. Esko, SARS-CoV-2 Infection Depends on Cellular Heparan Sulfate and ACE2. Cell (2020), doi:10.1016/j.cell.2020.09.033.

11. N. Volpi, R. J. Linhardt, High-performance liquid chromatography-mass spectrometry for mapping and sequencing glycosaminoglycan-derived oligosaccharides. Nature Protocols. 5, 993–1004 (2010).

12. N. Volpi, F. Galeotti, B. Yang, R. J. Linhardt, Analysis of glycosaminoglycan-derived, precolumn, 2-aminoacridone–labeled disaccharides with LC-fluorescence and LC-MS detection. Nature Protocols. 9, 541–558 (2014).

13. W. Wei, M. R. Niñonuevo, A. Sharma, L. M. Danan-Leon, J. A. Leary, A Comprehensive Compositional Analysis of Heparin/Heparan Sulfate-Derived Disaccharides from Human Serum. Anal. Chem. 83, 3703–3708 (2011).

14. H. Liu, Q. Liang, J. S. Sharp, Peracylation Coupled with Tandem Mass Spectrometry for Structural Sequencing of Sulfated Glycosaminoglycan Mixtures without Depolymerization. J. Am. Soc. Mass Spectrom. 31, 2061–2072 (2020).

15. X. Han, P. Sanderson, S. Nesheiwat, L. Lin, Y. Yu, F. Zhang, I. J. Amster, R. J. Linhardt, Structural analysis of urinary glycosaminoglycans from healthy human subjects. Glycobiology. 30, 143–151 (2020).

16. S. Tomatsu, T. Shimada, R. W. Mason, J. Kelly, W. A. LaMarr, E. Yasuda, Y. Shibata, H. Futatsumori, A. M. Montaño, S. Yamaguchi, Y. Suzuki, T. Orii, Assay for Glycosaminoglycans by Tandem Mass Spectrometry and its Applications. J Anal Bioanal Tech. 2014, 006 (2014).

17. W. Wei, R. L. Miller, J. A. Leary, Method development and analysis of free HS and HS in proteoglycans from pre-and postmenopausal women: evidence for biosynthetic pathway changes in sulfotransferase and sulfatase enzymes. Anal Chem. 85, 5917–5923 (2013).

18. X. Sun, L. Li, K. H. Overdier, L. A. Ammons, I. S. Douglas, C. C. Burlew, F. Zhang, E. P. Schmidt, L. Chi, R. J. Linhardt, Analysis of Total Human Urinary Glycosaminoglycan Disaccharides by Liquid Chromatography–Tandem Mass Spectrometry. Anal Chem. 87, 6220–6227 (2015).

19. Y.-H. Chen, Y. Narimatsu, T. M. Clausen, C. Gomes, R. Karlsson, C. Steentoft, C. B. Spliid, T. Gustavsson, A. Salanti, A. Persson, A. Malmström, D. Willén, U. Ellervik, E. P. Bennett, Y. Mao, H. Clausen, Z. Yang, The GAGOme: a cell-based library of displayed glycosaminoglycans. Nature Methods. 15, 881–888 (2018).

20. F. Maccari, F. Galeotti, V. Mantovani, L. Zampini, L. Padella, L. Rigon, D. Concolino, A. Fiumara, E. Pascale, A. Pittalà, T. Galeazzi, C. Monachesi, R. L. Marchesiello, G. Coppa, O. Gabrielli, N. Volpi, Composition and structure of glycosaminoglycans in DBS from 2-3-day-old newborns for the diagnosis of mucopolysaccharidosis. Anal Biochem. 557, 34–41 (2018).

21. F. Maccari, F. Galeotti, L. Zampini, L. Padella, R. Tomanin, D. Concolino, A. Fiumara, T. Galeazzi, G. Coppa, O. Gabrielli, N. Volpi, Total and single species of uronic acid-bearing glycosaminoglycans in urine of newborns of 2-3days of age for early diagnosis application. Clin Chim Acta. 463, 67–72 (2016).

22. J. C. Silva, M. S. Carvalho, X. Han, K. Xia, P. E. Mikael, J. M. S. Cabral, F. C. Ferreira, R. J. Linhardt, Compositional and structural analysis of glycosaminoglycans in cell-derived extracellular matrices. Glycoconj J. 36, 141–154 (2019).

23. B. Yang, Y. Chang, A. M. Weyers, E. Sterner, R. J. Linhardt, Disaccharide analysis of glycosaminoglycan mixtures by ultra-high-performance liquid chromatography-mass spectrometry. J Chromatogr A. 1225, 91–98 (2012).

24. N. K. Karamanos, Extracellular matrix: key structural and functional meshwork in health and disease. The FEBS Journal. 286, 2826–2829 (2019).

25. E. Huet, C. Jaroz, H. Q. Nguyen, Y. Belkacemi, A. de la Taille, V. Stavrinides, H. Whitaker, Stroma in normal and cancer wound healing. The FEBS Journal. 286, 2909–2920 (2019).

26. J. Wei, M. Hu, K. Huang, S. Lin, H. Du, Roles of Proteoglycans and Glycosaminoglycans in Cancer Development and Progression. International Journal of Molecular Sciences. 21, 5983 (2020).

27. M. Hu, Y. Lan, A. Lu, X. Ma, L. Zhang, in Progress in Molecular Biology and Translational Science (Elsevier, 2019; https://linkinghub.elsevier.com/retrieve/pii/S1877117318301509), xvol. 162, pp. 1–24.

28. L. Xu, L. Tang, L. Zhang, Proteoglycans as miscommunication biomarkers for cancer diagnosis. Prog Mol Biol Transl Sci. 162, 59–92 (2019).

29. F. Gatto, M. Maruzzo, C. Magro, U. Basso, J. Nielsen, Prognostic Value of Plasma and Urine Glycosaminoglycan Scores in Clear Cell Renal Cell Carcinoma. Front. Oncol. 6 (2016), doi:10.3389/fonc.2016.00253.

