## Supplemental Information for "Analytical Performance of a Standardized Kit for Mass Spectrometry-based Measurements of Human Glycosaminoglycans"

### SUPPLEMENTARY MATERIAL

#### Supplementary Methods

**Linearity of the Calibrators**. We prepared calibration standards for each disaccharide in triplicates, according to the kit instructions. In short, we prepared nine levels of calibration by a serial dilution from the highest level (Calibrator A). We reported the nominal concentration of the disaccharide at each calibration level in Table S1. We injected each of the replicates for UHPLC-MS/MS analysis in duplicates. After the UHPLC-MS/MS acquisition, we integrated all peaks with a signal-to-noise (S/N) ratio greater than 10. We then calculated the average peak area, standard deviation, and coefficient of variation (CV) for each level. We considered acceptable any calibration level with a CV < 25%, except for the lowest level where we deemed acceptable CV < 30%.

We established the calibration curve by performing a least-square regression between peak areas and the corresponding nominal concentrations. We then used the best fit curve to back-calculate the concentration at each calibration level. We discarded the measurements where the back-calculated concentration deviated more than 30 % from the nominal concentration and fitted a new regression iteratively.

**Selectivity and Specificity of the Calibrators.** We prepared three blank samples by processing Milli-Q water according to the kit instructions for the preparation of the calibration curve. Next, we injected the samples in UHPLC-MS/MS and we inspected the presence of a peak for each disaccharide. We quantified the apparent peak areas in the blank sample at the expected retention time for that disaccharide. Next, we compared the peak area to the lower limit of quantification (LLoQ) for that disaccharide. We considered the selectivity and acceptable if the area was lower than 20% LLoQ.

**Accuracy and Precision in the Calibrators.** We created a set of three standard GAG solutions at three different concentrations (low, medium, and high). We prepared the standard GAG solution at the "high" level by mixing the highest level of the calibrator sample for all disaccharides in milli-Q water. Next, we serially diluted the sample at the "high" level to "medium" and "low" levels (1 : 0.50 : 0.25 v/v) using Milli-Q water. We chose the dilution ratios to ensure that each disaccharide concentration was within the above-established linearity range. We performed the accuracy test over two consecutive days. We reported the accuracy for each disaccharide as the percentage difference between the nominal concentration and the measured concentration divided by the nominal concentration. Next, we estimated the precision of the calibration curve from the data generated in the accuracy test. In short, we calculated the precision as CV% on each day for each disaccharide. For a given disaccharide, we considered the accuracy acceptable if the difference was lower than 25% (30% at the lowest level). Similarly, we considered precision acceptable if the CV was lower than 25% (30% at the lowest level).

**Carry-over.** We prepared two blank samples according to the kit instructions. We assessed the carry-over by injecting blank samples after the acquisition of the sample corresponding to the highest level of calibration for all disaccharides. We inspected the presence of the peak for each disaccharide and quantified peak areas in the blank sample at the expected retention time for that disaccharide. We then compared the peak area to the LLoQ for that disaccharide. We consider carry-over negligible in cases where the peak area was lower than 20% LLoQ.

**Disaccharide Stability in the Autosampler.** We prepared the third level of calibrator in singleton according to the kit instructions by a serial dilution from the highest level (Calibrator A) for each disaccharide. We stored the sample in the autosampler at 10° C for the duration of the experiment. We quantified the peak area of each disaccharide daily until Day 8 and then on Day 14. We computed the percent change in peak area at each time point relative to the peak area at the initial time point for a given disaccharide. We considered the disaccharide stability in the autosampler acceptable if the change did not exceed 30% in at least 85% (e.g., 6 of 7 days) of consecutive time points including the last time point, which we then assumed to be the last day of stability for that disaccharide.

**Recovery.** We created five replicates of a set of three standard GAG solutions at three different concentrations (low, medium, and high), as described in “Accuracy and Precision”. We then prepared a “proxy urine” pool by mixing urine collected from healthy donors. Next, we depleted GAGs from the proxy urine pool by recovering the filtrate resulting from ultracentrifugation (14000 *g* at 9 °C for 60 minutes) in a filter provided in the kit. We spiked the standard GAG solution into the proxy urine at the beginning of sample preparation (before filtration) or immediately after filtration during sample preparation. We computed the recovery for each disaccharide as the ratio (in percentage) of the disaccharide concentration in the proxy urine sample spiked before filtration versus after filtration (assumed as reference concentration). We considered recoveries that deviated less than 25% from the reference concentration as acceptable.

**Matrix effect.** We created a set of two standard GAG solutions at two different concentrations (low and high), as described in “Accuracy and Precision”. We then prepared “proxy urine” from 6 healthy donors as described in “Recovery”. Immediately after filtration during the sample preparation, we spiked the standard GAG solution in each proxy urine sample. In parallel, we prepared a reference sample in triplicates by spiking the standard GAG solution in milli-Q water. We computed the matrix effect for each disaccharide as the ratio (in percentage) of the disaccharide concentration in each proxy urine sample versus the milli-Q water reference samples. We had no pre-specified acceptance criteria for matrix effects.

**Accuracy and Precision in Native Samples.** We prepared fresh proxy urine samples according to the kit instructions every day of the experiment. To account for day-to-day and operator variability, two operators repeated the experiment independently over three days. We prepared five replicates of a set of three standard GAG solutions at three different concentration levels (low, medium, and high), as described in "Accuracy and Precision", and spiked each replicate in a proxy urine sample. We determined the accuracy as the percentage difference between the nominal concentration of each disaccharide at a given level and the measured concentration divided by the nominal concentration. We used the same data to calculate the precision of the calibration curve, in terms of CV%, for each disaccharide. For a given disaccharide, we considered the accuracy acceptable if the difference was lower than 30% (35% at the lowest level). Similarly, we considered precision acceptable if the CV was lower than 25% (30% at the lowest level).

**Disaccharide Stability in Native Samples.** We prepared proxy urine samples as described in "Accuracy and Precision". Then, we prepared duplicates of a set of two standard GAG solutions at two different concentration levels (low and high), as described in "Accuracy and Precision", and spiked each replicate in a proxy urine sample. We stored proxy urine samples at -20°C throughout the experiment and subject to the same thaw/freeze cycle. We prepared an aliquot of each sample in triplicate according to the kit instructions on Day 1 and Day 14. For each GAG concentration level (low and high), we computed the percentage difference of each disaccharide concentration on Day 14 and Day 1 with respect to the corresponding nominal concentration. We considered disaccharide stability in native samples acceptable on Day 14 if the difference did not exceed 30%.

**Selectivity and Specificity in Native Samples.** We created proxy urine samples as described in "Recovery" and prepared them according to the kit instructions (without spiking of standard GAG solutions). We inspected the presence of peaks and eventually quantified peak areas in the proxy urine sample at the expected retention time for each disaccharide. We compared the peak area to the LLoQ for that disaccharide. We considered selectivity and specificity acceptable in cases where the area was lower than 20% LLoQ for that disaccharide.

**Linearity in Native Samples.** We created triplicates of a set of nine standard GAG solutions at nine different concentration levels, by serial dilution from the highest level as described in "Accuracy and Precision". We spiked each standard replicate in a proxy urine pooled sample. We performed a least-square linear regression between the peak area at each level and the corresponding concentration for each disaccharide. We considered the linearity acceptable if the coefficient of determination ($R^2$) was > 95%.

**Intra-laboratory Precision.** This analysis was conducted in an external laboratory, which followed Good Laboratory Practices wherever applicable (Lablytica Life Sciences AB, Uppsala, Sweden). One operator, trained in the use of the kit, repeated the experiment 14 times with four QC samples included in the kit, using a second lot of the kits. Two of the four QC samples were made of synthetic urine spiked by two concentration levels of standard GAG solutions ("Synthetic urine (low)" and "Synthetic urine (high)"). The remaining two QC samples were native urine samples from two healthy donors at two concentration levels of GAGs ("Native urine (low)" and "Native urine (high)"). QC samples were processed independently for each experiment and injected in UHPLC-MS/MS in duplicates. For each disaccharide, we monitored the concentration in each QC sample across the independent experiments. We computed the CV of each disaccharide from the mean and standard deviation across 14 runs (in percentage). We considered precision acceptable in cases where CV was less than 25% in "high" samples and less than 35% in "low" samples.

**Inter-laboratory Precision.** We prepared and analyzed a panel of 9 native urine samples from healthy donors independently in the reference laboratory according to kit instructions. The external laboratory prepared and analyzed the same samples in parallel, following the same procedure. The kits were part of a third lot of Elypta MIRAM$^{TM}$ Glycosaminoglycan Kit for Research Use-Only. We calculated estimates of the total CS concentration and 0s CS concentrations from the two laboratories and computed Pearson correlation coefficient $R$ and its $p$-value (permutation test). We considered the inter-laboratory precision acceptable if $R > 0.95$ for both GAG properties.

### Supplementary Figures

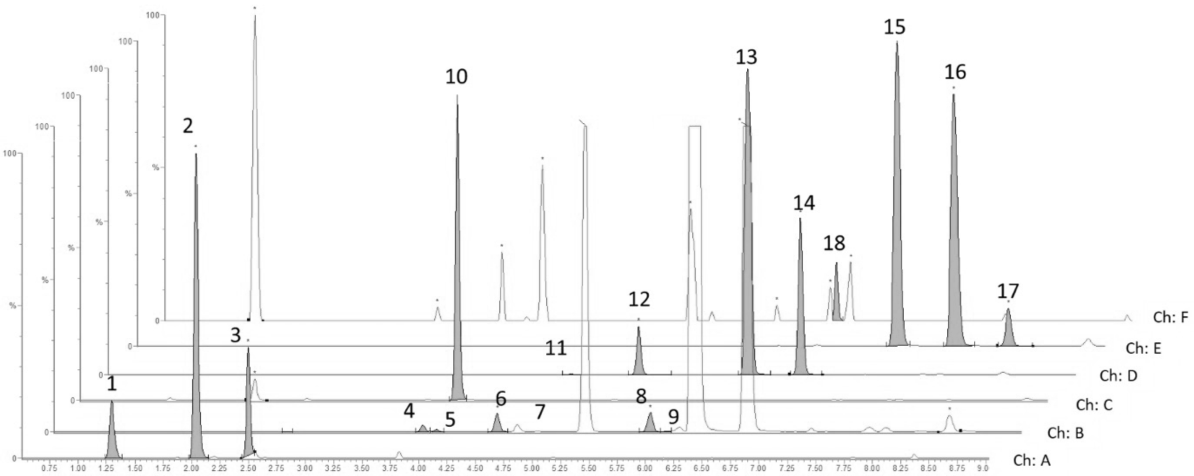


**Figure S1**. Representative chromatograms for a standard GAG solution consisting of a mixture of CS sodium (certified reference material), HS sodium salt (from bovine kidney) and sodium salt (from *Streptococcus equi*). Key - 1: Tris H; 2: Ns6s HS, 3: Ns2S HS, 4: Tris CS; 5: 2s4S CS; 6: 2s6s HS, 7: 2s6s CS, 8: 2s HS, 9: 2s CS, 10: Ns HS, 11: 4s6s CS, 12: 6s HS, 13: 4s CS, 14: 6s CS, 15: 0s HS, 16: HA, 17: 0s CS.

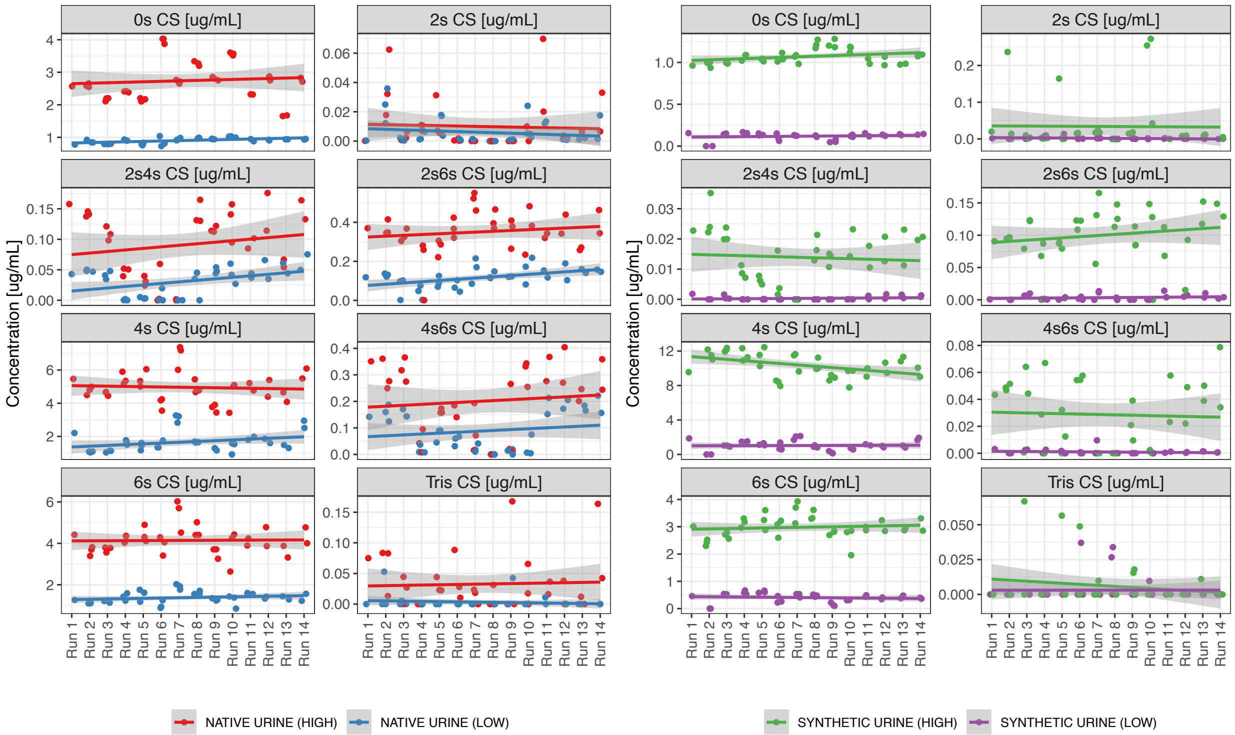


**Figure S2.** Measured CS GAG concentration (in µg mL$^{-1}$) in 4 QC samples (in duplicates) across 14 runs in an external laboratory. The line represents the least square regression for disaccharide concentration across runs (shaded area represents 95% confidence interval on the regressed mean).

### Supplementary Tables

**Table S1.** Nominal concentration of each disaccharide at each calibration level (A – I)

| **Disaccharide** | **Nominal concentration [µg/mL]** | | | | | | | | |
|---|---|---|---|---|---|---|---|---|---|
| | A | B | C | D | E | F | G | H | I |
| 0s CS | 5.330 | 3.020 | 1.712 | 0.970 | 0.550 | 0.311 | 0.176 | 0.100 | 0.057 |
| 4s CS | 42.870 | 24.293 | 13.766 | 7.801 | 4.420 | 2.505 | 1.419 | 0.804 | 0.456 |
| 6s CS | 11.150 | 6.318 | 3.580 | 2.029 | 1.150 | 0.651 | 0.369 | 0.209 | 0.119 |
| 4s6s CS | 0.310 | 0.176 | 0.100 | 0.056 | 0.032 | 0.018 | 0.010 | 0.006 | 0.003 |
| 2s4s CS | 0.110 | 0.062 | 0.035 | 0.020 | 0.011 | 0.006 | 0.004 | 0.002 | 0.001 |
| 2s6s CS | 0.600 | 0.340 | 0.193 | 0.109 | 0.062 | 0.035 | 0.020 | 0.011 | 0.006 |
| Tris CS | 0.040 | 0.023 | 0.013 | 0.007 | 0.004 | 0.002 | 0.001 | 0.001 | 0.000 |
| 2s CS | 0.040 | 0.023 | 0.013 | 0.007 | 0.004 | 0.002 | 0.001 | 0.001 | 0.000 |
| HA | 60.000 | 34.000 | 19.267 | 10.918 | 6.187 | 3.506 | 1.987 | 1.126 | 0.638 |
| Tris HS | 0.430 | 0.244 | 0.138 | 0.078 | 0.044 | 0.025 | 0.014 | 0.008 | 0.005 |
| Ns2s HS | 1.440 | 0.816 | 0.462 | 0.262 | 0.148 | 0.084 | 0.048 | 0.027 | 0.015 |
| 2s6s HS | 0.080 | 0.045 | 0.026 | 0.015 | 0.008 | 0.005 | 0.003 | 0.002 | 0.001 |
| Ns6s HS | 2.030 | 1.150 | 0.652 | 0.369 | 0.209 | 0.119 | 0.067 | 0.038 | 0.022 |
| Ns HS | 8.200 | 4.647 | 2.633 | 1.492 | 0.846 | 0.479 | 0.272 | 0.154 | 0.087 |
| 0s HS | 46.270 | 26.220 | 14.858 | 8.419 | 4.771 | 2.704 | 1.532 | 0.868 | 0.492 |
| 2s HS | 0.060 | 0.034 | 0.019 | 0.011 | 0.006 | 0.004 | 0.002 | 0.001 | 0.001 |
| 6s HS | 2.660 | 1.507 | 0.854 | 0.484 | 0.274 | 0.155 | 0.088 | 0.050 | 0.028 |

**Table S2.** Stability calculated as changes in disaccharide peak areas over 14 days in the autosampler at 10°C. Acceptable values are marked in bold. Key: ND – not detected.

| **Disaccharide** | **Day 0** | **Day 1** | **Day 2** | **Day 3** | **Day 4** | **Day 6** | **Day 7** | **Day 8** | **Day 14** |
|---|---|---|---|---|---|---|---|---|---|
| 0s CS | **100%** | **92%** | **89%** | **125%** | **124%** | **106%** | 142% | 142% | 164% |
| 4s CS | **100** | **85** | **78** | **106** | **103** | **90** | **103** | **104** | **111** |
| 6s CS | **100** | **85** | **75** | **127** | **115** | **89** | 147 | 148 | 191 |
| 4s6s CS | ND | ND | ND | ND | ND | ND | ND | ND | ND |
| 2s4s CS | **100** | **0** | **66** | **118** | **89** | **98** | **103** | **114** | 195 |
| 2s6s CS | **100** | **102** | **0** | **141** | **116** | **109** | 148 | 177 | 198 |
| Tris CS | ND | ND | ND | ND | ND | ND | ND | ND | ND |
| 2s CS | **100** | **78** | **95** | **110** | **123** | **106** | **129** | 154 | 159 |
| HA | **100** | **92** | **92** | 140 | 135 | **118** | 162 | 161 | 186 |
| Tris HS | ND | ND | ND | ND | ND | ND | ND | ND | ND |
| Ns2s HS | **100** | 0 | **125** | 8 | 142 | 192 | 6 | 272 | 392 |
| 2s6s HS | ND | ND | ND | ND | ND | ND | ND | ND | ND |
| Ns6s HS | **100** | **90** | **85** | **90** | **118** | **70** | **111** | **128** | 138 |
| Ns HS | **100** | **83** | **76** | **116** | **107** | **88** | **128** | **129** | 164 |
| 0s HS | **100** | **90** | **87** | **127** | **124** | **107** | 148 | 147 | 158 |
| 2s HS | **100** | 0 | **95** | **110** | **123** | **106** | **129** | 154 | 159 |
| 6s HS | ND | ND | ND | ND | ND | ND | ND | ND | ND |

**Table S3.** Recovery of each disaccharide in standard GAG solutions at three levels of concentration after filtration during sample preparation. Acceptable values are marked in bold. Key: ND – not detected.

| **Disaccharide** | **Low** | **Medium** | **High** |
|---|---:|---:|---:|
| 0s CS | **103%** | **103%** | **100%** |
| 4s CS | **102** | **102** | **96** |
| 6s CS | **104** | **103** | **94** |
| 4s6s CS | **107** | **105** | **108** |
| 2s4s CS | **105** | **102** | **94** |
| 2s6s CS | **106** | **105** | **92** |
| Tris CS | ND | ND | ND |
| 2s CS | 49 | 64 | **82** |
| HA | 171 | **123** | 284 |
| Tris HS | **88** | **85** | 139 |
| Ns2s HS | **93** | **123** | **113** |
| 2s6s HS | ND | ND | ND |
| Ns6s HS | **95** | **101** | 145 |
| Ns HS | **120** | **108** | 135 |
| 0s HS | **120** | **105** | 158 |
| 2s HS | **106** | 163 | 8 |
| 6s HS | **119** | **115** | 294 |

**Table S4**. Matrix effect (%) on the disaccharides' concentration in proxy urine (PU, $N = 6$) versus Milli-Q water samples spiked with two different concentration levels of a standard GAG solution. No acceptable values were pre-specified for matrix effects. Key: ND – not detected.

| **Disaccharide** | **PU1** | | **PU2** | | **PU3** | | **PU4** | | **PU5** | | **PU6** | |
|---|---|---|---|---|---|---|---|---|---|---|---|---|
| | L | H | L | H | L | H | L | H | L | H | L | H |
| **0s CS** | 97 | 74 | 94 | 86 | 100 | 65 | 98 | 69 | 88 | 78 | 95 | 77 |
| **4s CS** | 83 | 82 | 83 | 96 | 87 | 83 | 82 | 87 | 65 | 94 | 68 | 91 |
| **6s CS** | 93 | 63 | 93 | 73 | 95 | 58 | 98 | 62 | 84 | 70 | 91 | 64 |
| **4s6s CS** | 114 | 72 | 92 | 80 | 105 | 65 | 116 | 78 | 90 | 83 | 109 | 71 |
| **2s4s CS** | ND | 61 | ND | 58 | ND | 59 | ND | 61 | ND | 68 | ND | 70 |
| **2s6s CS** | 100 | 62 | 78 | 67 | 96 | 60 | 100 | 69 | 80 | 77 | 87 | 67 |
| **Tris CS** | ND | ND | ND | ND | ND | ND | ND | ND | ND | ND | ND | ND |
| **2s CS** | ND | 40 | ND | 42 | ND | 31 | ND | 44 | ND | 37 | ND | 58 |
| **HA** | 90 | 65 | 16 | 14 | 20 | 4 | 48 | 5 | 138 | 14 | 100 | 22 |
| **Tris HS** | ND | 53 | ND | 47 | ND | 27 | ND | 51 | ND | 87 | ND | 64 |
| **Ns2s HS** | ND | 68 | ND | 56 | ND | 34 | ND | 45 | ND | 79 | ND | 41 |
| **2s6s HS** | ND | ND | ND | ND | ND | ND | ND | ND | ND | ND | ND | ND |
| **Ns6s HS** | 97 | 82 | 34 | 51 | 85 | 55 | 91 | 23 | 101 | 45 | 119 | 37 |
| **Ns HS** | 134 | 74 | 33 | 27 | 104 | 35 | 160 | 42 | 227 | 44 | 173 | 37 |
| **0s HS** | 177 | 69 | 82 | 70 | 169 | 86 | 221 | 68 | 264 | 38 | 181 | 43 |
| **2s HS** | ND | ND | ND | ND | ND | ND | ND | ND | ND | ND | ND | ND |
| **6s HS** | 134 | 59 | 41 | 36 | 54 | 10 | 89 | 10 | 184 | 23 | 164 | 35 |

**Table S5.** Stability of disaccharide in proxy urine samples (in terms of deviation % from nominal concentration) at two concentration levels of a standard GAG solution on Day 1 and Day 14 while stored at -20°C. Note that stability for HA and HS disaccharides could not be reliably estimated in this experiment and it was omitted. Acceptable values marked in bold. Key: ND – not detected.

| **Disaccharide** | **Low** | | **High** | |
|---|---|---|---|---|
| | Day 1 | Day 14 | Day 1 | Day 14 |
| 0s CS | **9%** | **25%** | **15%** | 37% |
| 4s CS | **16** | **5** | **3** | 4 |
| 6s CS | **-17** | **-3** | **5** | **10** |
| 4s6s CS | **6** | **27** | 31 | 33 |
| 2s4s CS | ND | ND | 13 | 29 |
| 2s6s CS | **-8** | **17** | **-7** | **19** |
| Tris CS | ND | ND | ND | ND |
| 2s CS | ND | ND | ND | ND |

**Table S6**. Coefficient of variation (%) for each disaccharide in 4 QC samples included in the kit across 14 independent runs in an external laboratory. In bold, acceptable CV% at high (<25%) or low (<35%) concentrations.

| Disaccharide | Synthetic (HIGH) | Synthetic (LOW) | Native (HIGH) | Native (LOW) |
|---|---|---|---|---|
| Total CS | **8%** | **33%** | **11%** | **19%** |
| 4s CS | **11** | 42 | **16** | **35** |
| 6s CS | **11** | **26** | **12** | **16** |
| 0s CS | **8** | **15** | **22** | **8** |
| 2s6s CS | 27 | 83 | **22** | **31** |
| 4s6s CS | 65 | 109 | 55 | 76 |
| 2s4s CS | 50 | 132 | 56 | 65 |
| Tris CS | 147 | 263 | 83 | 207 |
| 2s CS | 167 | 109 | 141 | 126 |
| Total HS | 27 | **20** | 64 | **35** |
| 0s HS | 33 | **22** | 54 | 39 |
| Ns HS | 31 | **23** | 58 | 38 |
| 6s HS | 28 | **26** | 65 | 46 |
| Ns6s HS | 41 | 103 | 117 | 71 |
| Ns2s HS | 75 | 132 | 160 | 171 |
| 2s6s HS | 48 | 78 | 138 | 125 |
| Tris HS | 206 | 361 | 205 | 246 |
| 2s HS | 56 | 67 | 123 | 74 |
| Total HA | **24** | **34** | 139% | 36% |
| Total GAG | **13** | **18** | **12** | **20** |
